# Dormancy dampens the microbial distance-decay relationship

**DOI:** 10.1101/717546

**Authors:** KJ Locey, ME Muscarella, ML Larsen, SR Bray, SE Jones, JT Lennon

**Affiliations:** Department of Biology, Indiana University; School of Science, Technology, Engineering and Math, Diné College; Department of Plant Biology, University of Illinois; Wilfrid Laurier University; Department of Biology, Transylvania University; Department of Biological Sciences, University of Notre Dame

**Keywords:** dormancy, seed banks, distance-decay, biogeography, environmental filtering, microbial ecology

## Abstract

Much of Earth’s biodiversity has the capacity to engage in dormancy whereby individuals enter a reversible state of reduced metabolic activity. By increasing resilience to unfavorable conditions, dormancy leads to the accumulation of “seed banks” that should diminish the influence of environmental filtering, while allowing passive dispersers to colonize new habitats. Although prevalent among single-celled organisms, evidence that dormancy influences patterns of microbial biodiversity and biogeography is lacking. We constructed geographical and environmental distance-decay relationships (DDRs) using 16S rRNA sequencing to characterize the total (DNA) and the active (RNA) bacterial communities in a regional survey of 49 forested ponds. As expected, the total community harbored greater diversity and exhibited weaker DDRs than the active portion of the community. These empirical observations were robust to different measures of community similarity and random resampling tests. Furthermore, findings from the field survey were reproduced by models that included aspects of dormancy along with the geographical coordinates and environmental characteristics of our study system. In addition to maintaining local diversity, our results support recent theoretical predictions that dormancy shapes geographical patterns of biodiversity.

## INTRODUCTION

Dormancy is a reversible state of reduced metabolic activity that is exhibited in various forms across the tree of life [1, 2]. Among plants, animals, and the microbial life that dominates our planet, many organisms contribute to the collection of dormant individuals known as a seed bank [2-4]. Seed banks are often viewed as the outcome of a bet-hedging strategy that allows populations to distribute reproductive effort among long-lived individuals over time [1, 5, 6]. By increasing survival in fluctuating environments, dormancy also reduces the probability of local extinctions and maintains diversity via the storage effect [2, 7-9]. In addition, dormancy can influence the commonness and rarity of species in a community while allowing propagules to tolerate conditions that would otherwise prohibit dispersal and colonization [2, 9-11]. Although crucial for understanding the local dynamics of populations and communities [12-14], dormancy is historically neglected when attempting to predict and explain patterns of biodiversity.

The influence of dormancy on general relationships of biodiversity and biogeography should perhaps be strongest among microbial communities. Microorganisms are well known to accumulate in seed banks by forming cysts, spores, and other morphological resting stages that are highly resilient to fluctuations in a wide range of environmental conditions. For example, recent estimates suggest there are more than 10^29^ endospores in the marine subsurface alone [15]. In addition, microorganisms can engage in less conspicuous forms of dormancy that involve the downregulation of metabolic activities, which allows individuals to extend their lifespan for prolonged periods of time [16]. As a result, microbial seed banks can account for the vast majority of individuals and biomass in some ecosystems [1, 15, 17, 18]. Historically, microorganisms have also been thought to have relatively unhindered capacities for dispersal, in part, through the influence that dormancy has on improving the odds of successful dispersal and colonization [9]. Through its influence on dispersal and environmental filtering, microbial dormancy is not only expected to increase the number of species in a given habitat (α-diversity), but also decrease the turnover of species (β-diversity) across similar habitats of a given landscape [9, 19].

One classical biogeographical pattern that is likely to be influenced by microbial dormancy is the distance decay relationship (DDR). The DDR characterizes how community similarity changes across increasing geographical distance [20], often reflecting the influence of dispersal limitation and spatially autocorrelated environmental conditions [21-24]. The DDR has attracted attention among microbial ecologists for testing predictions of island biogeography [25], for elucidating scale-dependency of beta-diversity [26], and for revealing the relative influence of geographical isolation versus environmental conditions [27-30]. Because it can decrease environmental filtering and dispersal limitation, microbial dormancy is predicted to flatten the slope of the DDR [19, 31]. For the same reasons, we expect dormancy to also decrease the similarity between communities that are found in close geographic and environmental proximity to one another, which would be reflected by the *y*-intercept of the DDR.

In the current study, we sample and analyze communities of bacteria from a regional distribution of small freshwater ponds. Using 16S rRNA sequencing, we constructed geographical and environmental DDRs by characterizing the compositional similarities among ponds using the total (DNA) and active (RNA) portions of the bacterial communities. Going beyond previous studies in several ways, we compare not only the slopes but also the *y*-intercepts of geographical and environmental DDRs for the total and active communities. Likewise, we base our analyses on multiple aspects of community similarity including commonness and rarity. To augment our field study, we constructed simulation-based models that encode our hypothesized mechanisms. We challenged the models to reproduce our empirical results, and evaluated whether greater environmental filtering, decreased death in dormancy, and increased dormant dispersal lead our models to more closely approximate our empirical results. In accordance with recent predictions, our findings support the hypothesis that patterns of microbial biogeography are weakened by seed banks, which may help explain discrepancies in macroecological patterns across domains of life.

## METHODS

### Field study

#### Study system and environmental data

Between 1950 and 1960, approximately 300 ponds were constructed as wildlife habitat in Brown County State Park (BCSP), Yellowwood State Forest (YSF), and Hoosier National Forest (HNF) in south central Indiana, USA (Fig. 1). Built along topographical ridgelines, the ponds are hydrologically disconnected from one another within the forested landscape. We identified a subset (*n* = 49) of ponds to sample based on their representation across watersheds and their accessibility, which was influenced by proximity to recreational trails, fire trails, or a logging road. Because surrounding vegetation can shade these small ponds and affect community structure [32], we quantified the percentage of visible sky above the center of every pond by capturing hemispherical images of the forest canopy. In addition to measuring other spatial and physical properties (elevation, depth, and diameter), we used a YSI 556 MPS multiprobe to quantify water temperature (°C), redox potential (ORP, mV), specific conductivity or (SpC, µS/cm), dissolved oxygen (DO, mg/L), total dissolved solids (TDS, g/L), and salinity (PSU, ppm). We then collected surface water samples (0.1 m) from the center of each pond using an extended pole to prevent sedimentary disturbance and transported them back to the laboratory for additional analyses. Specifically, we measured dissolved organic carbon (DOC) and dissolved organic nitrogen (DON) concentrations on 0.7-µm filtrates via high-temperature combustion and chemiluminescence with a Shimadzu TOC-V/TNM instrument. We measured water color (*a*_440_) as a surrogate for terrestrial-derived dissolved organic matter [33]. We quantified total phosphorus (TP) after persulfate digestion via spectrophotometric analysis using a molybdenum blue analysis [34]. Last, we estimated algal biomass by quantifying chlorophyll *a* with a Turner Designs fluorometer after cold-extracting biomass that was retained on 0.7-µm glass fiber filters in 95 % ethanol [35].

**Fig. 1.**
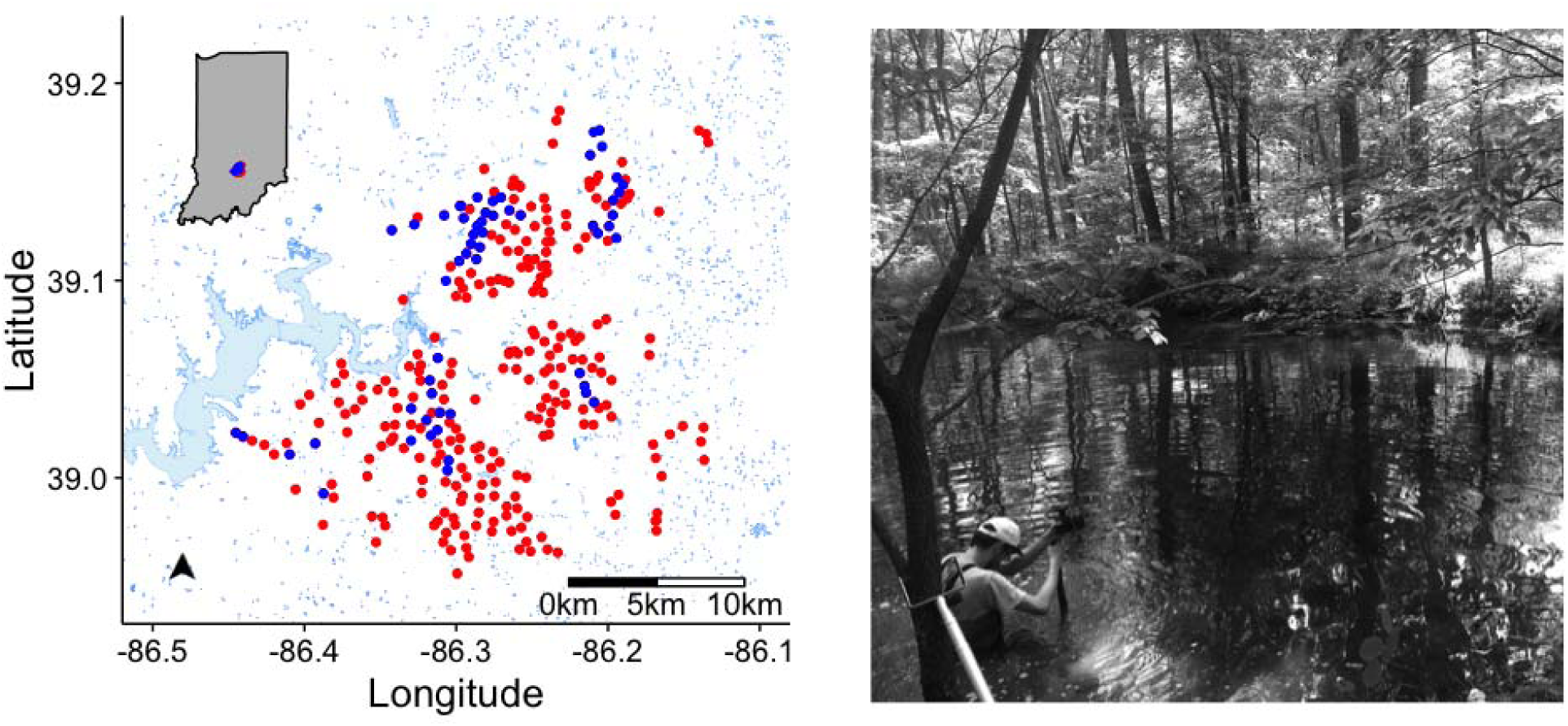
Left: A map of our study area depicting the locations of forested ponds constructed in south central Indiana, USA. For this study, we sampled 49 of the ponds (blue symbols); the remaining ponds were not sampled (red symbols). Surrounding surface water is depicted in light blue shapes. Right: A typical pond showing its relatively small size and surrounding vegetation.

#### Sampling and sequencing microbial communities

We characterized bacterial communities by sequencing 16S rRNA genes and transcripts from pond water samples. This approach has been previously used to analyze the composition of the active (RNA) and total (DNA) taxa in complex microbial samples [11, 36, 37] based on the following logic. Sequences recovered from the DNA pool can be derived from organisms that are dead, dormant, or active [38]. In contrast, ribosomal RNA, which is required for protein synthesis, has a short half-life and is only produced by metabolically active individuals [39, 40]. Thus, through the simultaneous analysis of 16S rRNA genes and transcripts, we can gain insight into how seed banks contribute to biogeographic patterns of microbial communities [1]. For each pond sample, we harvested planktonic microbial biomass by filtering surface-water onto a 47 mm 0.2 µm Supor Filter (Pall) using vacuum filtration. We then co-extracted nucleic acids from one half of the filter using the QIAGEN RNeasy PowerWater Kit and a DNA elution accessory kit. We eliminated DNA from RNA samples using the RTS DNase kit (QIAGEN) and synthesized first strand cDNA using the SuperScript III First Strand Synthesis Kit (Invitrogen) with random hexamer primers. Then, we amplified the V4 region of the 16S rRNA gene from the DNA and cDNA templates using barcoded 515F and 806R primers [41]. After purifying with the AmPure XP kit (Beckman), we quantified amplicons using the Quant-iT kit (Invitrogen) and then pooled libraries at approximately equal molar ratios (20 ng each). Following procedures outlined in greater detail elsewhere [38, 42], we sequenced the pooled library on the Illumina MiSeq platform using 250 × 250 base pair (bp) paired-end reads (Illumina Reagent Kit v2) at the Indiana University Center for Genomics and Bioinformatics Sequencing Facility. Raw sequence reads were assembled into contigs, quality-filtered to remove long sequences and sequences with ambiguous bases, and aligned to the Silva reference database (version 123). We used the UCHIME algorithm [43] to detect and remove chimeric sequences. Additionally, we removed mitochondrial, archaeal, and eukaryotic sequences following detection with the Ribosomal Data Base Project’s 16S rRNA reference sequences and taxonomy version 6 [44]. Finally, we split the sequences based on taxonomic class using the RDP taxonomy and binned them into operational taxonomic units (OTUs) based on 97 % sequence similarity. All initial sequence processing was completed using the software package mothur (version 1.32.1, [45]).

#### Quantifying distance-decay relationships (DDRs)

We constructed DDRs for both the active (RNA) and total (DNA) communities by calculating pair-wise community similarity among sites (i.e., ponds). Because community similarity is not an absolute measure and depends on aspects of the communities being compared (e.g., dominance, rarity, presence-only), we used three distance-based metrics that capture contrasting properties of similarity. First, we used Bray-Curtis similarity, a common metric that emphasizes the influence of abundant taxa:

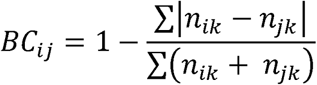

where *n*_*ik*_ is the abundance of taxon *k* at the *i*^*th*^ and *j*^*th*^ sites. Second, we calculated the Sørensen metric, which is a binary form of Bray-Curtis that accounts for presence of taxa in a sample, and hence, gives equal weight to rare and abundant taxa. Finally, we used the Canberra similarity metric, which places emphasis on the abundance of rare taxa:

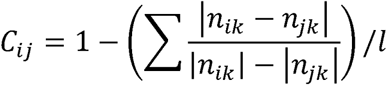

where *l* is the length of the community vector in the site-by-taxa matrix. For both the Bray-Curtis and Canberra measures, we used relative abundances to decrease the site-specific bias that occurs when total sample abundances of two sites greatly differ. We calculated each measure of community similarity using the *vegdist*() function in the ‘vegan’ R package [46]. We then calculated geographical and environmental distances among our sample ponds. For geographical distances, we used the ‘fossil’ R package [47]. For environmental distances, we used pair-wise differences in scores from the first axis of the principle components analysis (PCA), which was conducted on centered and rescaled environmental data using the ‘vegan’ R package [46]. Given the disparate ranges in environmental and geographical distance, we relativized distances to values between 0 and 1 by dividing each pair-wise distance by the maximum distance. This procedure also made environmental and geographical distance scale across the same range as community similarity, i.e., 0 to 1.

To test for the effect of dormancy on bacterial biogeography, we calculated the slopes and intercepts of environmental and geographical DDRs for the active and total communities using ordinary least-squares regressions [20, 29, 48]. We then tested whether differences in slopes and intercepts of DDRs were statistically significant using a permutation-based linear regression approach [20], which was implemented using the *diffslope*() function in the ‘simba’ R package. We also quantified DDRs using 5, 10, 15, 20, 25, 30, 35, and 45 ponds chosen at random to control for the influence of the samples size, and then repeated this 1,000 times to obtain mean values for each sample size. Additionally, we evaluated the effect of taxon richness on DDR parameters by randomly selecting different numbers of OTUs (1000, 4000, 8000, … 22,000) and rerunning our analyses. Because this process involved randomly selecting columns from the site-by-taxa matrices, we could not randomly select OTUs based on relative abundance or numbers of reads in individual samples (i.e., communities).

### Simulation modeling

#### Rationale

Empirical relationships like the DDR are often used to test ecological hypotheses but cannot, by themselves, demonstrate whether hypothesized mechanisms can reproduce observed relationships. This is further complicated by assumptions and idiosyncrasies that are inherent to empirical data. For example, studies that characterize microbial communities using high-throughput DNA sequencing implicitly assume that primer coverage, extraction efficiency, and amplification bias do not prevent molecular surveys from reflecting actual community structure[49]. There are also assumptions regarding the relationship between growth rate and rRNA content, when making inferences about the metabolic activity of microbial taxa using 16S ribosomal DNA and RNA sequencing [50, 51]. However, these caveats and limitations can, in part, be addressed by coupling empirical efforts with models that explicitly encode hypothesized mechanisms and that are independent of methodological assumptions. Therefore, we developed a simulation platform to test whether the environmental and geographical DDRs observed among ponds can be reproduced by focusing on the influence of environmental filtering, dormancy, and dispersal.

#### Overview

To evaluate the relative importance of dispersal and dormancy, we modeled population abundances within and among communities by simulating site-by-taxa matrices and using the same geographical coordinates, environmental PCA coordinates, and number of sites as in our field survey. By using the same magnitudes of richness and maximum abundances as in our field surveys, we avoided conflating the influence of environmental filtering, dormancy, and dispersal with general demographic differences between simulated and empirical data. As with species abundance models of community ecology, our models did not simulate across time steps but instead provide a “snapshot in time” [52]. Nevertheless, our approach yielded tens of thousands of models, each simulating several matrices having nearly one million elements each.

#### Simulating environmental filtering

To simulate environmental filtering, local abundances of dormant and active populations were determined by the match between site-specific values of the first PCA axis (PCA_1_) and a set of species-specific environmental optima. For simplicity, a single optimum was chosen at random for each species, by randomly selecting values within the range of scores from the first PCA axis (PCA_1_). For example, if the PCA_1_ scores ranged between −6 and 6, then a species was assigned a single optimum by drawing a uniform random number between −6 and 6. To simulate greater degrees of environmental filtering, we increased the range of randomly chosen species optima beyond the range of PCA_1_ scores. For example, a species could have a randomly chosen environmental optimum of −10 to 10. In this way, greater environmental filtering led to a greater number of species being poorly fit to a greater number of locations. The match (*m*_*ij*_) between the environmental optimum of the *i*^*th*^ taxon (*O*_*j*_) and the PCA_1_ score at the *j*^*th*^ site (*E*_*i*_) was calculated as follows:

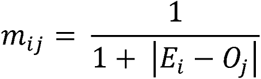

Values of *m*_*ij*_ ranged between 0.0 and 1.0, corresponding to a zero match and perfect match, respectively. For each species, a larger *m*_*ij*_ would result in greater active abundance while a smaller *m*_*ij*_ resulted in greater death in the active populations and larger dormant populations.

#### Simulating dormant and active abundances

Using a basic niche-based assumption, active populations of each species occurred at each location with a probability equal to *m*_*ij*_. To approximate the magnitude of the greatest active OTU abundances in our field-based survey, active populations were initiated with 50,000 individuals. We assumed that dormant populations were greatly influenced by stochasticity and were essentially invisible to environmental filtering [9, 53]. Thus, we initiated the occurrence of dormant populations with a binomial probability of 0.08, i.e., the average probability of occurrence for taxa within our empirical data, and assigned randomly chosen abundances ranging between 1 and 10,000. A fraction of each active population equal to 1 - *m*_*ij*_ was then assigned to its corresponding dormant subpopulation. Active populations were then decreased in proportion to their environmental mismatch (1 – *m*_*ij*_). Hence, occurrence, abundance, and death among active populations was determined by environmental filtering (via *m*_*ij*_). Dormant populations were decreased by a randomly chosen percentage (0 - 100), the value of which varied across simulations and allowed us to test for the influence of dormant death. Hence, death in dormancy was independent of environmental conditions while dormant abundances were partly influenced by environmental filtering and partly by process-based stochasticity, i.e., neutral processes.

#### Simulating dispersal

To simulate the effect of dispersal between dormant populations and between active populations, the number of immigrating individuals was determined by three conditions. Specifically, the number of immigrating propagules was influenced by the pair-wise distance (km) between sites (*d*_*ij*_), the local population sizes of the donor site (*n*_*i*_), and percentages (*p*_*active*_, *p*_*dormant*_) that were randomly chosen across simulations. Hence, the number of propagules received by an OTU at site *j* from the same OTU at site *i* was equal to 1/(1+*d*_*ij*_) * *n*_*i*_ * *p*.

#### Model runs and analysis

We ran 8·10^4^ simulations, each with randomly drawn parameter values that influenced environmental filtering, dormant death rate, and active and dormant dispersal. This random sampling of parameters was designed to explore the complete space of model behavior and compare the relative importance of dormancy, dispersal, and environmental filtering. When completed, we combined the simulated active and dormant site-by-species matrices to form the site-by-species matrix for the total community, and then analyzed our simulated communities in an identical manner to the data collected from our field survey. Specifically, for each model we estimated the slopes and y-intercepts of geographical and environmental DDRs for the active and total communities. To draw inferences about the effects of environmental filtering, dormancy, and dispersal on the biogeographic patterns of the active and dormant fractions, we calculated the percent error (100 · |simulated – empirical|/empirical) between the slopes and *y*-intercepts of field-based and simulation-based DDRs.

## RESULTS

### Field survey

#### Environmental and geographic features

In the 500 km^2^ study region, the sampled ponds were separated by an average distance of 9.7 km (± 6.15 SD), an average nearest neighbor distance of 2.18 km (± 3.70 SD), and an average furthest neighbor distance of 24.9 km (± 5.10 SD). Despite being constructed by the same team of people over a relatively short period of time, the morphological characteristics of the ponds were quite different. There was a four-fold range in pond diameter (range = 7 - 32.5 m) and a six-fold range in maximum depth (range = 0.3 - 2.0 m). In addition, environmental conditions varied substantially among the sampled ponds, often with >100 % difference between minimum and maximum values (Table 1). The first two axes of the PCA explained 40.3 % and 12.5 % of the variation, respectively, in environmental conditions across sampled ponds. Along the first PCA axis, larger ponds with a more open canopy and elevated oxygen levels separated from smaller ponds with higher concentrations of nutrients, and suspended solids (Fig. S1). These multivariate observations are consistent with other strong relationships among environmental variables including a positive correlation between specific conductivity and dissolved organic nitrogen (Fig. S2). Importantly, geographical and environmental distances among ponds were unrelated (*p* > 0.5) (Fig. S3). Because of this decoupling, we were able to directly compare environmental DDRs and geographic DDRs for bacterial communities without any additional statistical treatment.

**Table 1.**
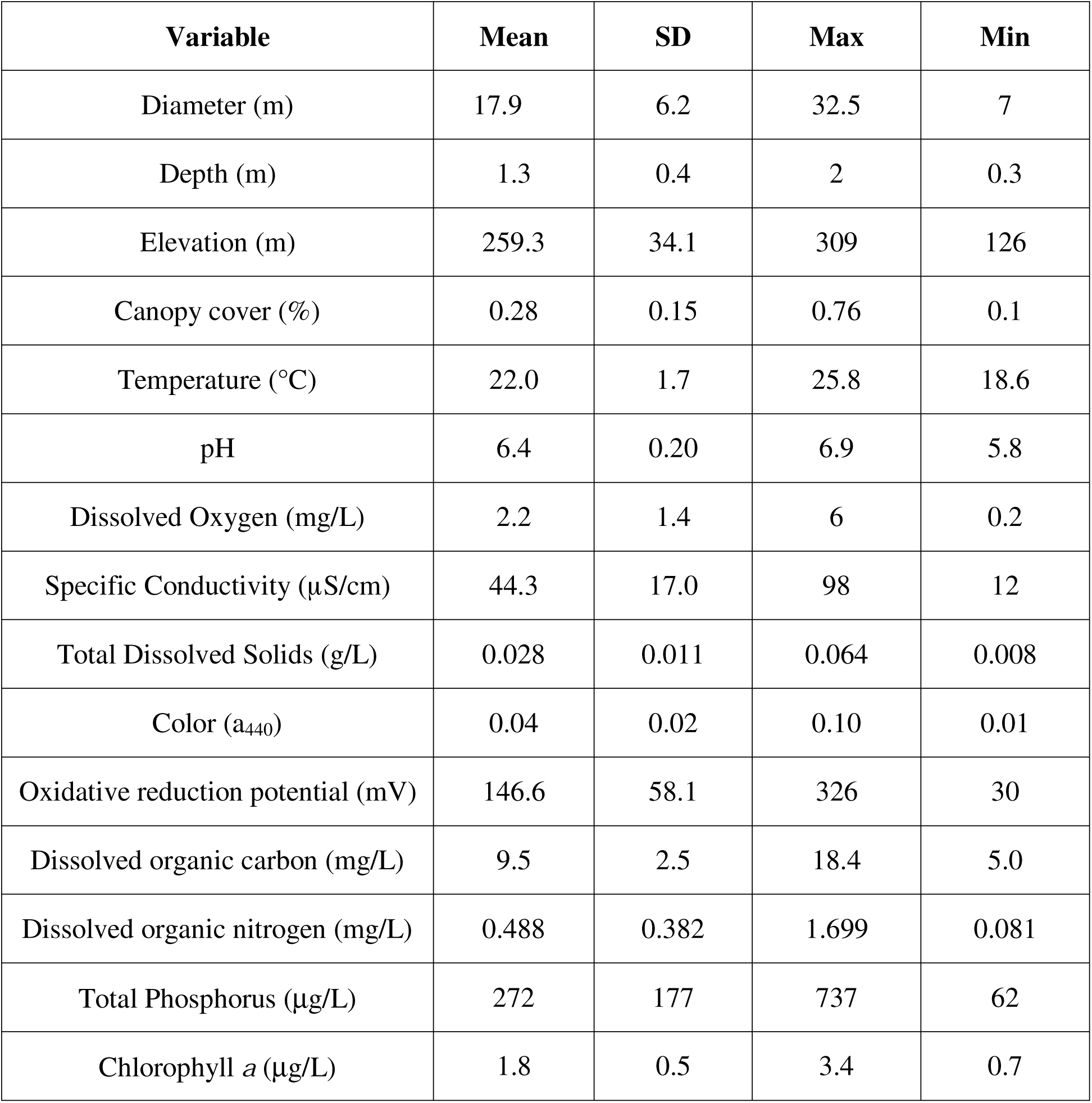
Descriptive statistics for 16 environmental variables measured from 49 forested ponds in southern Indiana, USA.

#### Diversity of active and total communities

As expected, active communities (RNA) were less diverse than the total (DNA) community. Across all ponds, we observed 21,568 OTUs among our active communities and 27,446 OTUs among our total communities. Within a pond, we observed on average 1,854 (± 586 SD) active OTUs and 2,083 (± 733 SD) total OTUs, which corresponded to a site-specific difference of 229 (± 507 SD) taxa.

#### Distance decay of bacterial communities

As predicted, DDRs for the active (RNA) community were characterized by steeper slopes and greater *y*-intercepts than those of the total (DNA) community (Fig. 2, Fig S4-S5, Tables 2-3). Through random resampling, we showed that these differences were independent of the number of sites (Fig. S6) and the number of OTUs (Fig. S7). While our findings were generally robust to community similarity metrics, the distinction between DDR parameters for the active and total communities was greatest when rare taxa were emphasized using either the Canberra or Sørensen index (Fig 2, Figs S4-S5, Tables 2-3). However, DDR slopes were steepest and intercepts were greatest when similarity was measured via Bray-Curtis. Bringing these two latter results together, it appears that community similarity was driven largely by dominant OTUs while differences between the active and total communities were driven by rare OTUs (Fig 2, Fig S4-S5, Tables 2-3). We documented this contrast between the active and total community for both environmental and geographical DDRs (Fig. 2, Tables 2-3, Table S2). However, environmental DDRs had steeper slopes and greater y-intercepts than geographical DDRs. By randomly permuting communities (i.e., rows) from the site-by-taxa matrices, we found that the slopes and *y*-intercepts of environmental DDRs differed from a null expectation. In contrast, slopes of geographical DDRs only differed from a null expectation when community similarity emphasized dominant OTUs, i.e., Bray-Curtis (Fig 2, Fig S4-S5).

**Table 2.** Slopes and summary statistics associated with the distance decay relationships (DDRs) based on environmental distance, geographical distance, and three different similarity metrics (Bray-Curtis, Sørensen, Canberra). Slopes were derived from simple linear regression models; *P*-values correspond with tests to determine whether DDR slopes from the active bacterial community (RNA-based) are different from the total bacterial community (DNA-based).

| Metric | DDR type | Community | Slope | % difference | *P*-value |
| --- | --- | --- | --- | --- | --- |
| Bray-Curtis | Environmental | Active | -0.326 | 25.2 | 0.001 |
|  |  | Total | -0.253 |  |  |
|  | Geographical | Active | -0.182 | 31.8 | 0.015 |
|  |  | Total | -0.132 |  |  |
| Sørensen | Environmental | Active | -0.099 | 57.1 | 0.001 |
|  |  | Total | -0.055 |  |  |
|  | Geographical | Active | -0.052 | 47.6 | 0.002 |
|  |  | Total | -0.032 |  |  |
| Canberra | Environmental | Active | -0.053 | 58.5 | 0.001 |
|  |  | Total | -0.029 |  |  |
|  | Geographical | Active | -0.028 | 54.5 | 0.002 |
|  |  | Total | -0.016 |  |  |

**Table 3.** Intercepts and summary statistics associated with the distance decay relationships (DDRs) based on environmental distance, geographical distance, and three different similarity metrics (Bray-Curtis, Sørensen, Canberra). Intercepts were derived from simple linear regression models; *P*-values correspond with tests to determine whether DDR intercepts from the active bacterial community (RNA-based) are different from the total bacterial community (DNA-based).

| Metric | DDR type | Community | Intercept | % difference | *p*-value |
| --- | --- | --- | --- | --- | --- |
| Bray-Curtis | Environmental | Active | 0.434 | 0.69 | 0.332 |
|  |  | Total | 0.437 |  |  |
|  | Geographical | Active | 0.422 | 0.94 | 0.343 |
|  |  | Total | 0.426 |  |  |
| Sørensen | Environmental | Active | 0.319 | 21.5 | 0.001 |
|  |  | Total | 0.257 |  |  |
|  | Geographical | Active | 0.314 | 20.7 | 0.001 |
|  |  | Total | 0.255 |  |  |
| Canberra | Environmental | Active | 0.106 | 26.7 | 0.001 |
|  |  | Total | 0.081 |  |  |
|  | Geographical | Active | 0.103 | 25.1 | 0.001 |
|  |  | Total | 0.080 |  |  |

**Figure 2.**
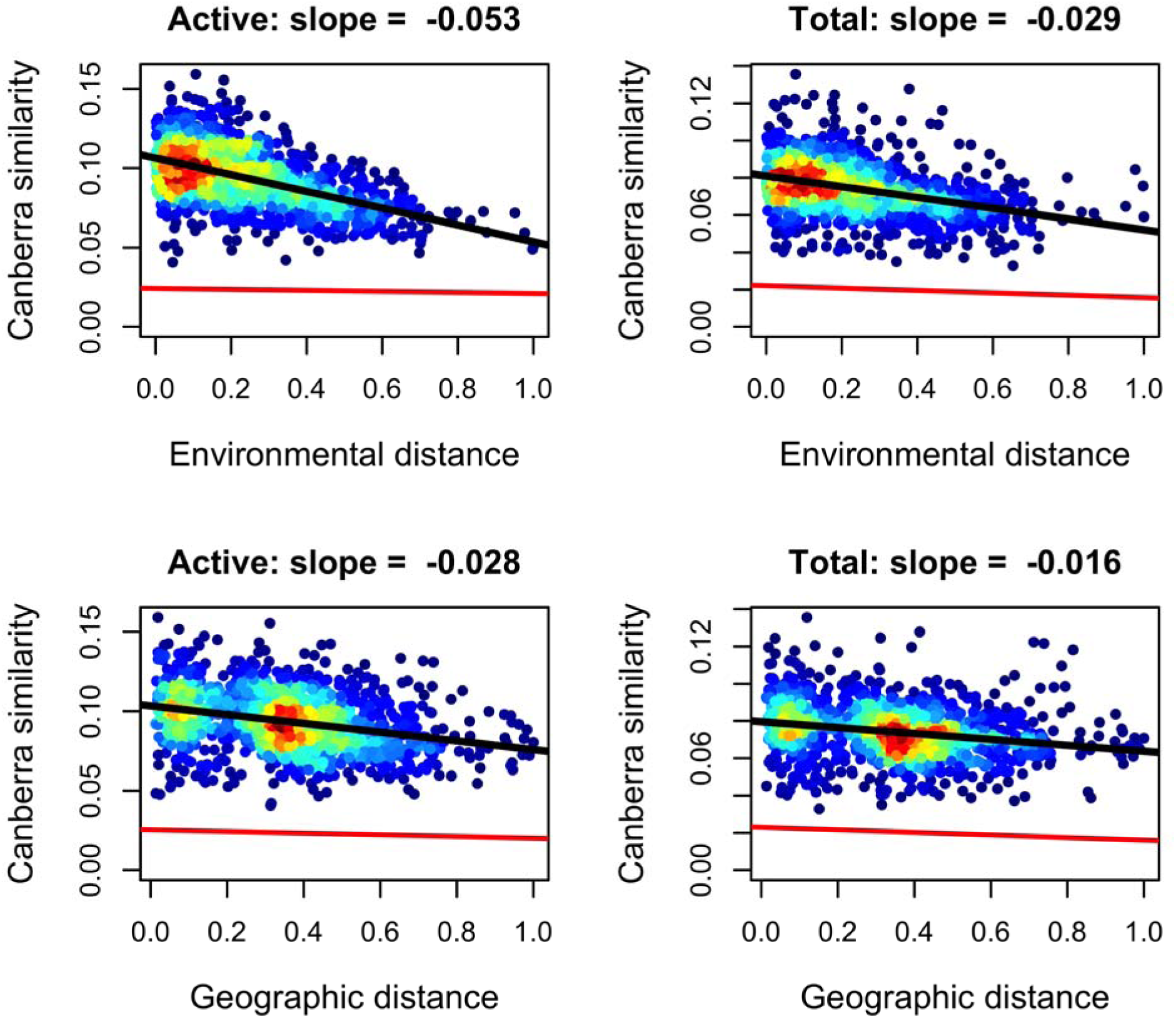
Distance-decay relationships (DDRs) for active (RNA-based) and total (DNA-based) bacterial communities sampled from forested ponds. Using Canberra metric of community similarity, which emphasizes rare taxa, we found that environmental DDRs (top row) were 58.5 % steeper for the active community than for the total community (*P* = 0.001). Geographical DDRs (bottom row) were 54.5 % steeper for the active community than for the total community (*P* = 0.001). Warmer colors represent a greater density of data points. Analyses based on Bray-Curtis distance and Sørensen’s distance produced qualitatively similar results (see Supplementary Materials). The red line is the DDR resulting from a null model whereby communities were randomly assigned to different locations.

### Simulation modeling

Models, whether simulation-based or not, are rarely ever challenged to reproduce more than one or two aspects of empirical data (e.g., the slope of a single DDR). However, nearly 7,900 of our simulations were capable of reproducing >50 aspects of our empirical results (Table S1). For example, across varying degrees of environmental filtering and dormant death, we found that: environmental and geographical DDRs from our simulations were statistically significant and less than 0 (*p* < 0.05), intercepts were greater than 0, slopes of environmental DDRs were steeper than those of geographical DDRs, DDRs for the active community were characterized by steeper slopes and greater *y*-intercepts than those of the total community, and each of these results held irrespective of the similarity metric (see Table 2).

Among the subset of simulations that reproduced these and other aspects of our empirical DDRs, those with the highest levels of environmental filtering and lowest levels of dormant death came within 10 % of our empirical DDR slopes (Fig. 3, Figs. S11-S12). In contrast, dispersal between sites had little-to-no effect on how closely a given simulation approximated the slopes of our empirical DDRs (Fig. 3, Figs. S11-S12). These latter results support the lack of an effect of geographic distance and dispersal among ponds in driving our empirical DDR findings. In contrast to the notion that “everything is everywhere, but the environment selects”, our empirical and simulation-based findings do not reflect cosmopolitan distributions nor unhindered dispersal. Instead, the simulations point to the overwhelming influence of environmental filtering and decreased rates of death due to dormancy, i.e., “…the environments selects.”

**Fig. 3.**
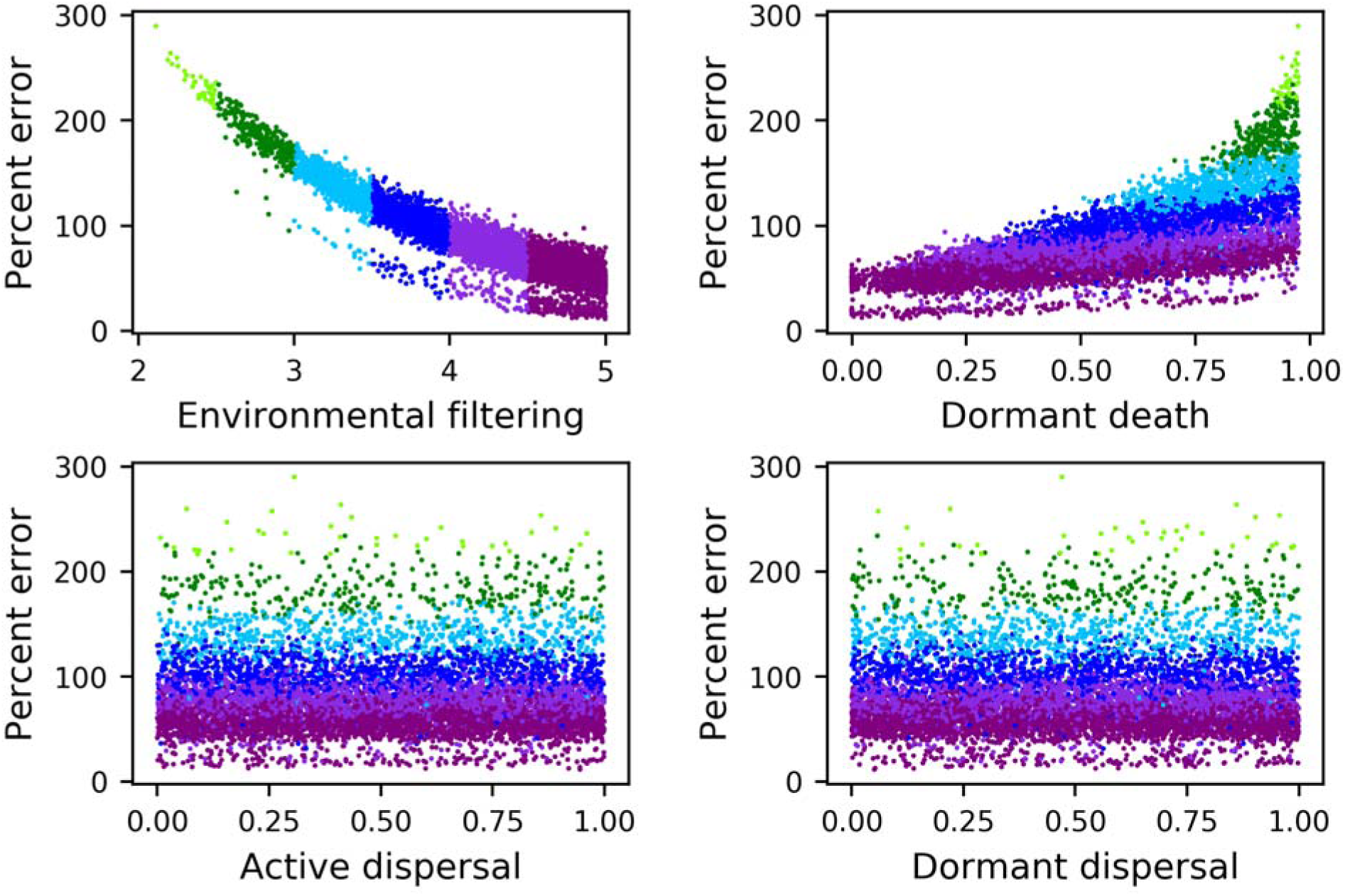
Average percent error between slopes of DDRs from the field survey and slopes of DDRs from simulated communities versus randomly drawn values of model parameters. Community similarity is based on the Canberra metric. Each point represents the result of a single model and is colored via the value of environmental filtering. Models with greater degrees of environmental filtering and lesser degrees of dormant death more closely approximated empirical DDRs. Dispersal among sites had no effect on the degree to which simulated DDRs approximated empirical DDRs. Similar results were produced with Bray-Curtis and Sørensen’s distances (see Supplementary Materials).

## DISCUSSION

In nature, organisms are commonly challenged by environmental conditions that are suboptimal for growth and reproduction. Throughout the tree of life, dormancy has evolved as a life-history strategy that helps ensure the persistence of populations through time. While numerous studies have demonstrated how dormancy maintains local diversity, few studies have explored how seed banks affect spatial patterns of biodiversity, including biogeographic relationships. In this study, we characterized bacterial diversity from a regional-scale survey of ponds in a forested landscape using an RNA- and DNA-based sequencing approach with the goal of constructing distance decay relationships (DDRs) for the active and total portions of the community, respectively. We augmented our field surveys with *in silico* simulations to test whether dormancy and environmental filtering could reproduce various aspects of our empirical results and approximate the parameters (slopes and intercepts) of our environmental and geographical DDRs. Our findings suggest that dormancy dampened patterns of biogeography within our study system and that these DDRs were largely driven by environmental filtering and resilience of seed banks to local conditions.

### No evidence that dispersal influenced DDRs

Unhindered dispersal, whether facilitated by dormancy or not, should weaken biogeographical patterns in microbial systems. Our field surveys revealed little-to-no support for the influence of geographical distance on DDRs. Similarly, our simulations revealed no support for the effect of dispersal on DDRs. In several cases, slopes of geographical DDRs did not differ from our null model and dispersal between sites was unnecessary for our simulation models to reproduce our empirical findings. However, these findings should not be interpreted as evidence for the absence of dispersal in our study system. While the spatial extent and average distance among sample sites was not nearly as low as some studies [19], the effect of spatially correlated dispersal on passively dispersing organisms should be greatest at scales where the mechanisms responsible for passive dispersal (e.g., oceanic and air currents, long-distance migrations) have a low chance of being spatially random [54, 55].

### Environmental filtering, dormancy, and their interaction

Our field surveys and simulations suggest that environmental filtering and dormancy have strong effects on community similarity. While environmental DDRs for the total community were significant, DDRs for the active community were stronger (Fig S4-S5, Tables 2-3). However, rather than having independent effects, interactions between environmental filtering and dormancy likely shape the structure of microbial communities. The dormant community does not just harbor taxa that immigrate into a suboptimal environment. Rather, the composition of dormant and active communities should reflect the repeated transitions between activity and dormancy that are inevitably driven by local-to-regional environmental changes. Because the death of dormant cells should be less influenced by environmental conditions than the death of active cells, DDRs for the active community should, as we predicted and demonstrated, be stronger than DDRs for the total community.

### The influence of dormancy on commonness and rarity

Commonness and rarity have been increasingly studied in microbial ecology, often with consideration given to the potential influence of dormancy [56-58]. In our study, we found that environmental and geographical DDRs were strongest (steeper slopes, greater *y*-intercepts) when community similarity emphasized dominant taxa. However, differences in DDR slopes and intercepts between the active and total communities were greatest when similarity emphasized rare taxa or ignored abundance altogether. Together, these findings suggest that compositional similarity across distance is more strongly driven by dominant taxa, but also that the influence of dormancy in dampening DDRs is manifested in the compositional differences that dormancy produces among rare taxa. There is at least one straightforward reason for these observations. Specifically, the abundance and occurrence of rare (i.e., low abundance) taxa are more likely than dominant taxa to be affected by random variation. This effect would explain the lower *y*-intercepts and shallower slopes of DDRs based on Canberra distance that were observed not only in our empirical data but also in our simulations which lacked biological or methodological assumptions associated with molecular surveys of diversity. In sum, dormancy appears to have an effect on the commonness and rarity of microbial taxa with implications for local- and regional-scale patterns of diversity [11, 56-58].

### Greater insight through modeling

In our study, we capitalized on the consistency of empirical and model-based results for making strong inference. Active communities in our field surveys had steeper DDRs and greater intercepts for both environmental and geographical DDRs. These findings were robust to different community similarity metrics and random resampling of the data. Our empirical results were then reproduced by thousands of simulations that were based on basic ecological assumptions, while being free of methodological caveats associated with RNA-DNA comparisons. Consequently, in absence of direct observations on the dormant community, the combination of strong empirical results and models that consistently and closely reproduce those empirical results can be particularly powerful in studying microbial phenomena that are difficult to directly observe, such as, dormancy and microbial mortality, but also heterogeneity in active metabolism, ribosomal turnover, dispersal routes, and dispersal distances.

### Greater insights through documenting dormancy

Large portions of microbial abundance and diversity can be found within dormant seed banks[1, 15]. The direct observation of seed banks is a significant challenge for microbial ecology and for the greater understanding of microbial diversity and biogeography. However, given the potential of developing technologies to directly survey microbial seed banks, future research might examine the degree to which environmental shifts drive compensatory changes in the composition of the active and dormant communities and the degree to which various forms of dormancy (e.g., spores vs. persistor cells) influence microbial biodiversity and biogeography. Additionally, while theoretical studies of microbial dormancy have assumed that entire taxa go dormant or become active [9], the degree to which portions of a given taxon’s abundance tends to be found in both the seed bank and the active community remains an unresolved area of active research [59].

#### Conclusion

The study of microbial biogeography has matured from a century-old hypothesis of unhindered dispersal and cosmopolitan distributions to a body of literature that supports many of the basic tenets of biogeography and that has given increasing consideration to the nuance of spatial scale [29, 55, 60]. In particular, dispersal limitation and environmental filtering are among the most general forces that shape ecological relationships and biogeographical patterns, and the effects of these forces should neither be independent of each other, independent of the influence of dormancy, nor operate equally across all scales. Dormancy is likely an important but historically overlooked process that influences biogeography and biodiversity [2]. This may explain why patterns of microbial biogeography often appear dampened in comparison to what is reported for plants and animal communities. Because of the inherent challenges in documenting dispersal, dormancy, environmental filtering, microbial ecologists should seek multiple lines of converging evidence, along with creative combinations of empirical study and modeling when considering how different processes potentially give rise to similar outcomes.

## ACKNOWLEDGEMENTS

We acknowledge J Eagleman and SA Cortwright for providing historical information about the forested ponds of southern Indiana, field assistance from S Widney and B Nowinski, laboratory assistance from BK Lehmkuhl and S Kranke, and financial support from the National Science Foundation DEB-1442246 (JTL, KJL, SEJ) and the US Army Research Office Grant W911NF-14-1-0411 (JTL). Data and code used in this manuscript can be found on GitHub (https://github.com/LennonLab/DormancyDecay) and NCBI (BioProject PRJNA554375).

