## Supplementary File for "Dormancy dampens the microbial distance-decay relationship"

**Supplemental figures and table**

**Figure S1**. Principle component analysis (PCA) conducted on 17 environmental variables.

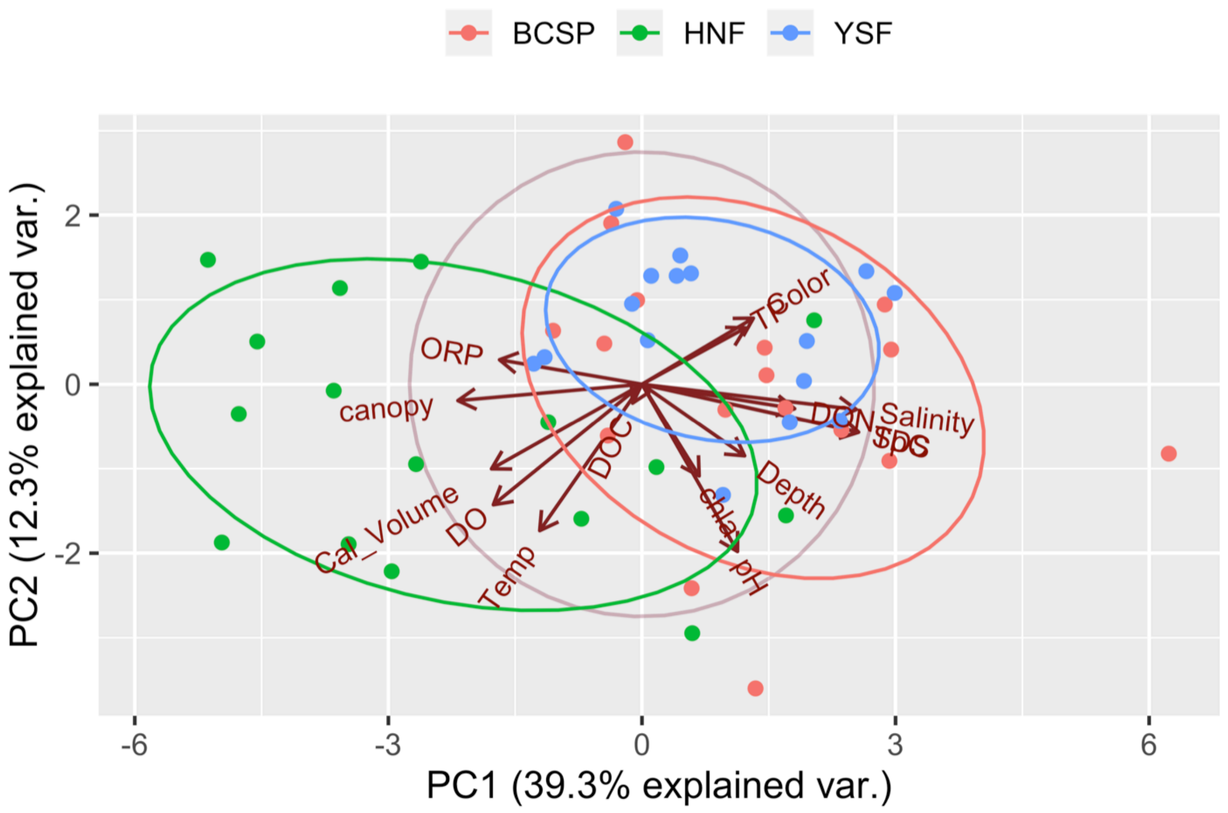

**Figure S2.** Correlation plot for environmental variables from 49 forested ponds.

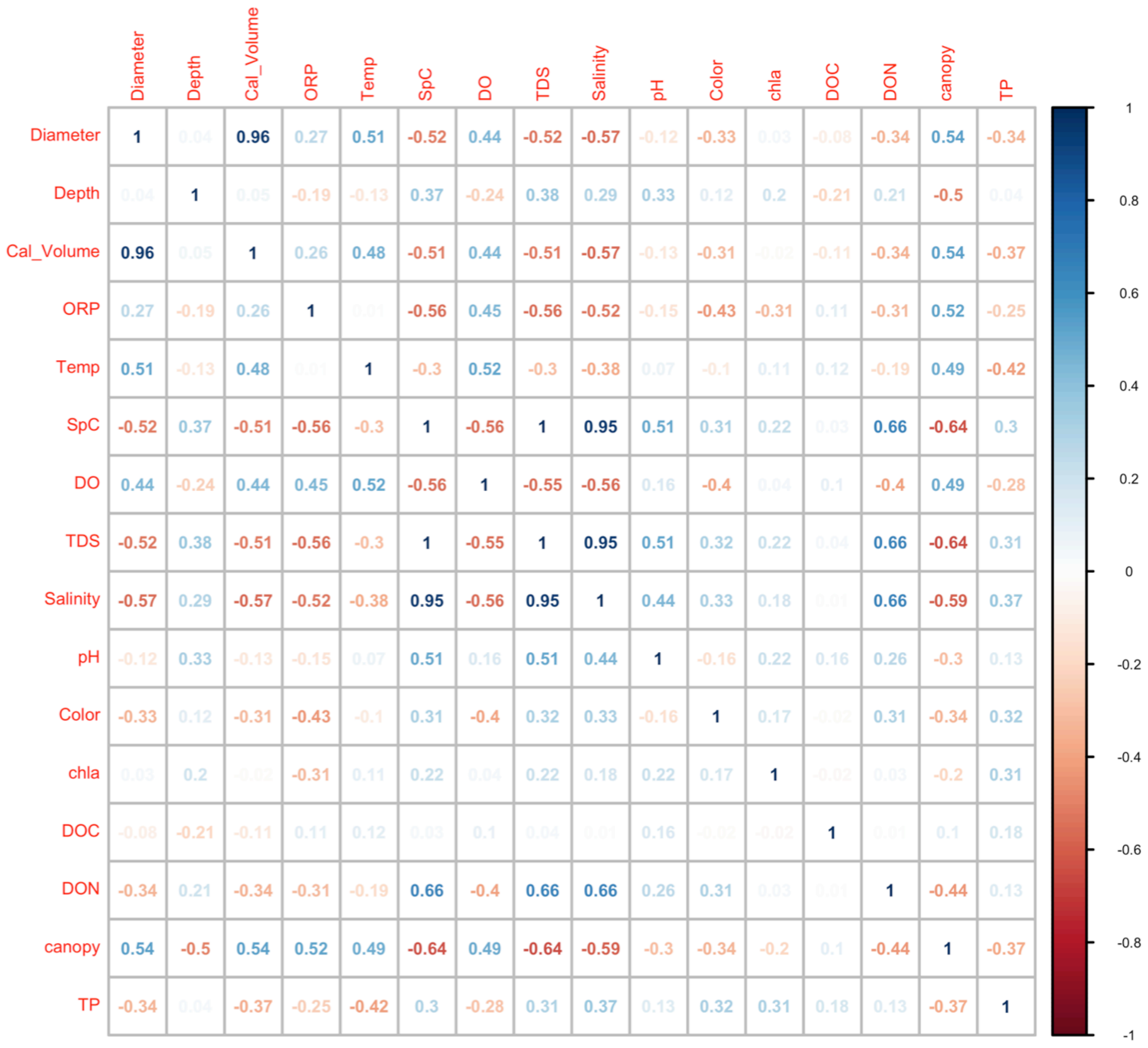

**Figure S3.** Geographical and environmental distances among ponds were unrelated ($P > 0.5$) (Fig. S3).

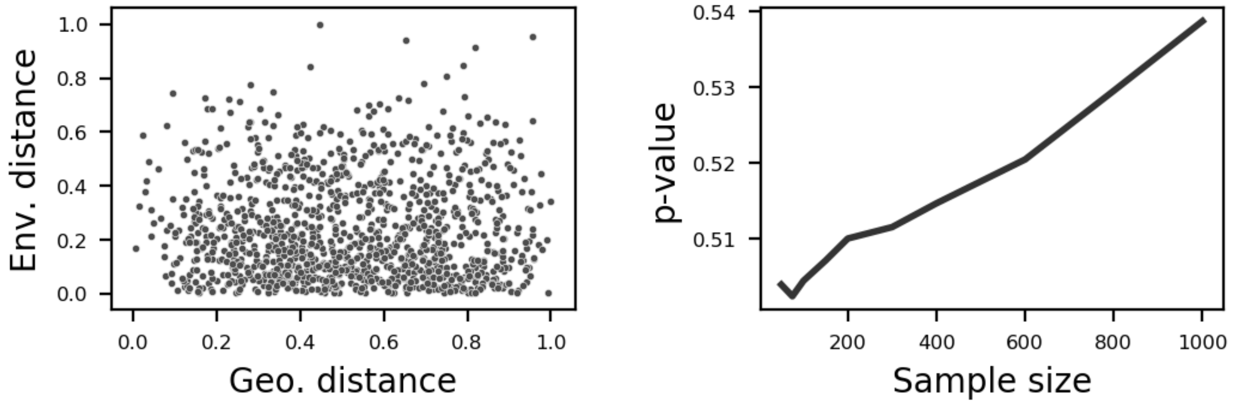

**Figure S4.** Results for distance decay relationships (DDR) for Bray-Curtis similarity. Environmental DDRs for the active community (RNA) and for the total community (DNA) differed by 25.2% ($p = 0.002$). Geographical DDRs for the active community (RNA) and for the total community (DNA) differed by 31.8 % ($P = 0.016$). The red line is the DDR resulting from a null model whereby communities were randomly assigned to different locations.

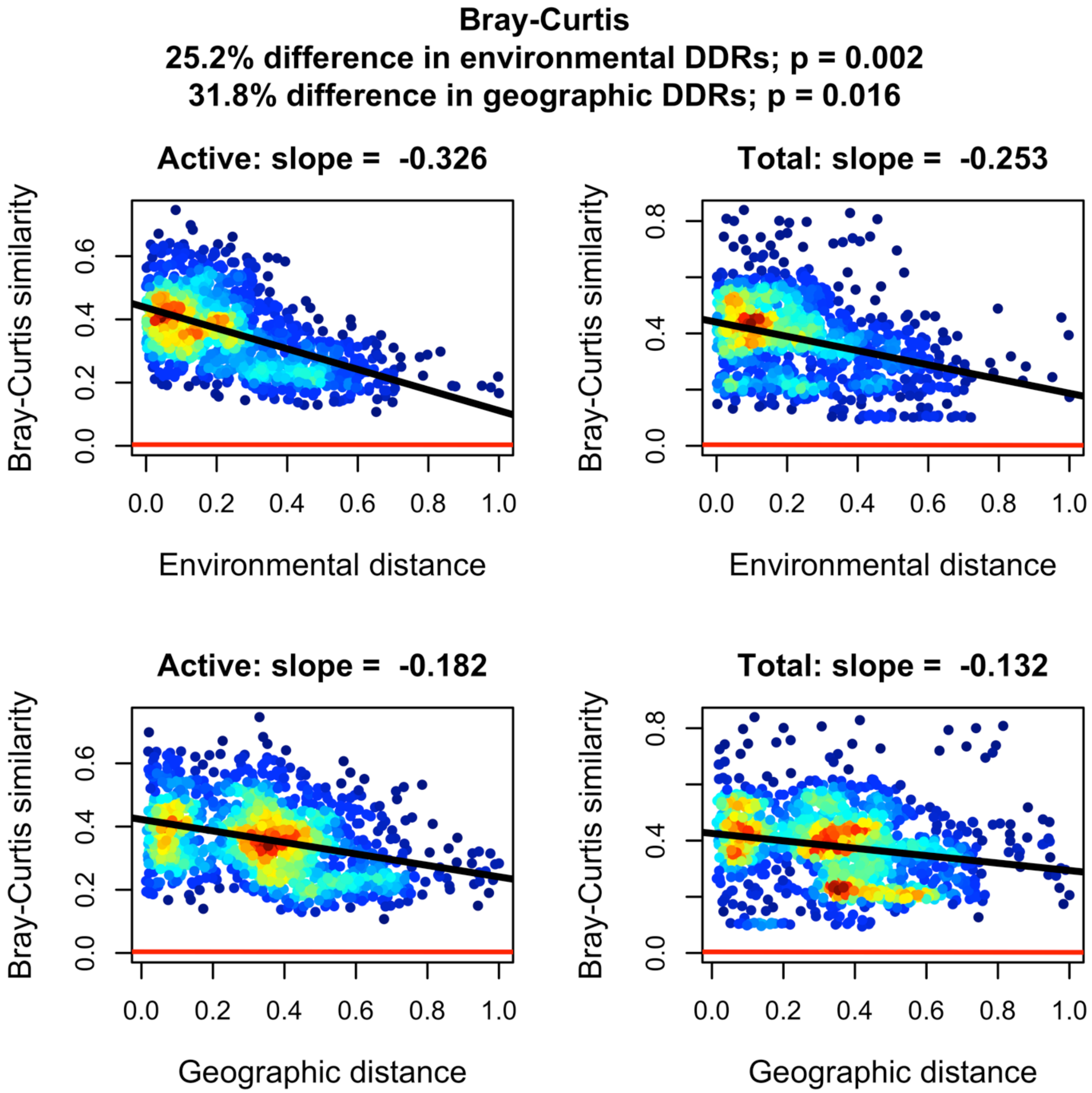

**Figure S5.** Results for distance decay relationships (DDR) for Sørensen's similarity. Environmental DDRs for the active community (RNA) and for the total community (DNA) differed by 57.1 % ($P = 0.001$). Geographical DDRs for the active community (RNA) and for the total community (DNA) differed by 47.6 % ($P = 0.003$). The red line is the DDR resulting from a null model whereby communities were randomly assigned to different locations.

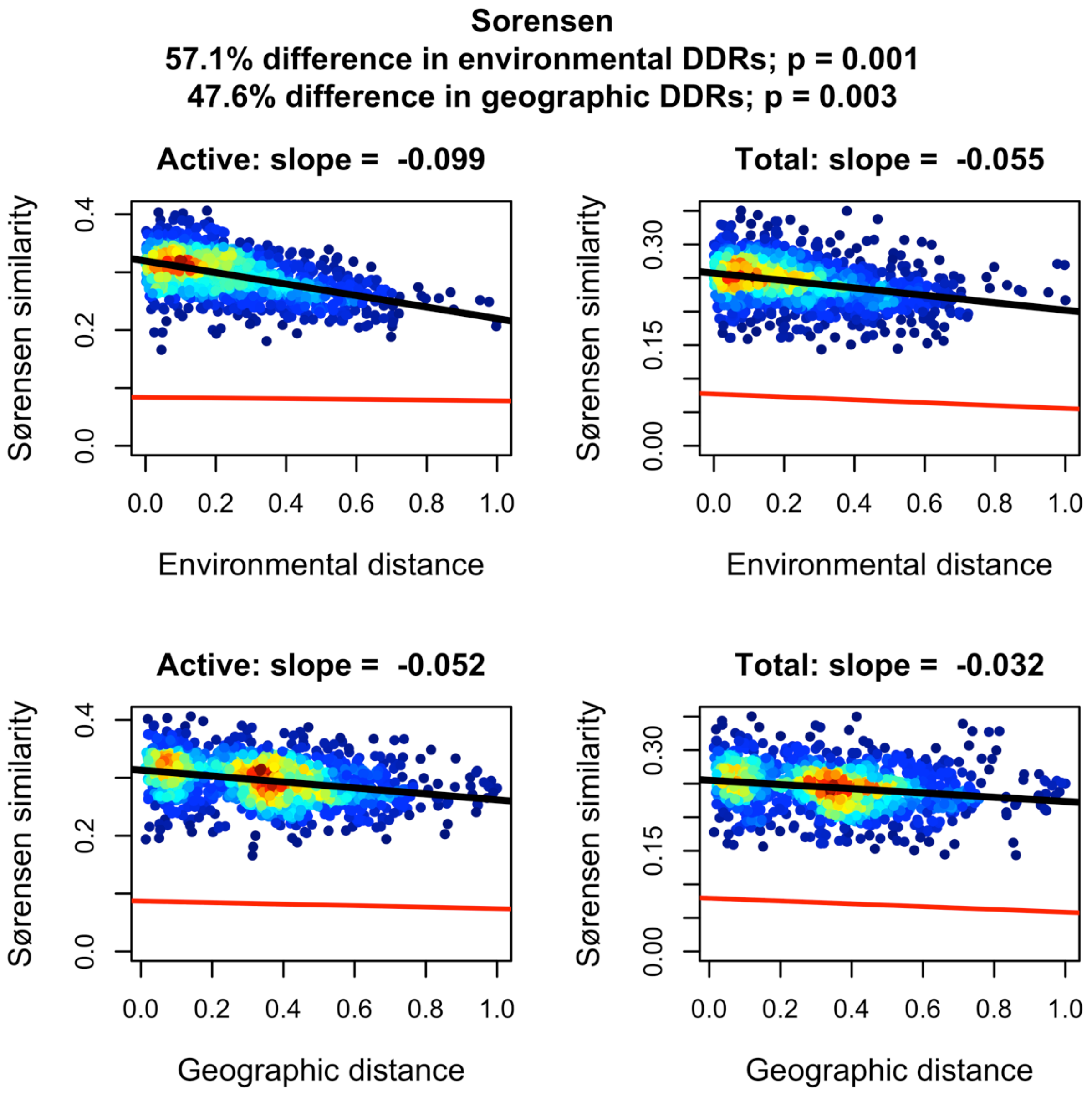

**Figure S6.** Slopes of environmental and geographical DDRs were steeper for the active community than for the total community regardless of the number of randomly chosen sample sites. Standard error bars and average values (points) were generated from 100 random resamplings.

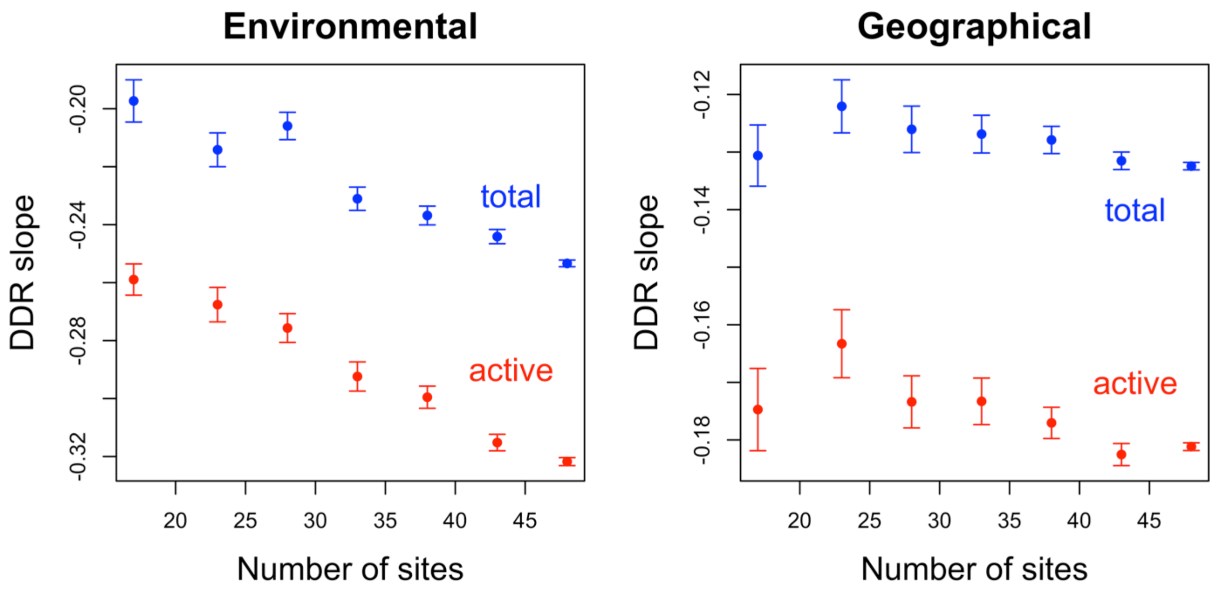

**Figure S7.** Slopes of environmental and geographical DDRs were steeper for the active community than for the total community regardless of the number of randomly chosen operational taxonomic units (OTUs). Columns (OTUs) were randomly chosen from site-by-taxa matrices. Abundances were then relativized. Standard error bars and average values (points) were generated from 100 random resamplings.

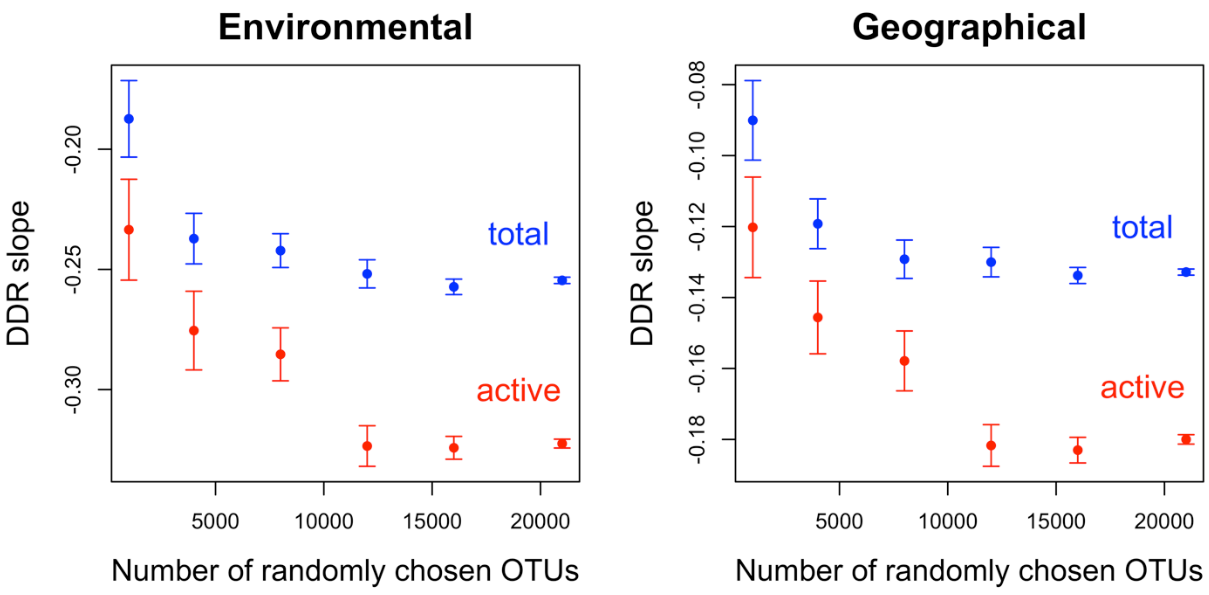

**Figure S8.** Results for distance decay relationships (DDR) for Sørensen's similarity. For the active community (RNA), environmental DDRs and geographical DDRs differed by 62.3 % ($P$ = 0.001). For the total community (DNA), environmental DDRs and geographical DDRs differed by 52.9 % ($P$ = 0.001). The red line is the DDR resulting from a null model whereby communities were randomly assigned to different locations.

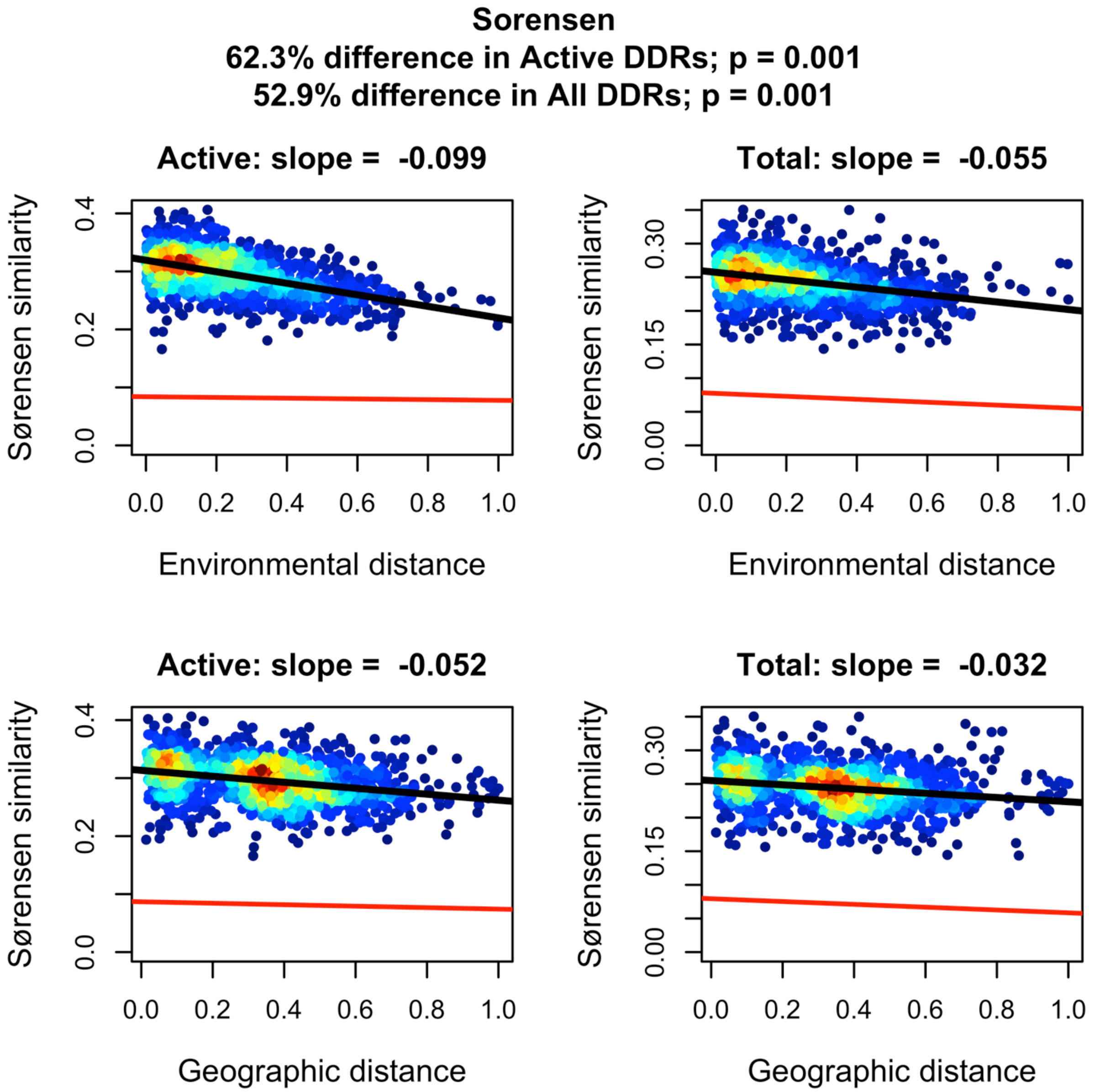

**Figure S9.** Results for distance decay relationships (DDR) for Bray-Curtis similarity. For the active community (RNA), environmental DDRs and geographical DDRs differed by 56.7 % ($P$ = 0.001). For the total community (DNA), environmental DDRs and geographical DDRs differed by 62.9 % ($P$ = 0.001). The red line is the DDR resulting from a null model whereby communities were randomly assigned to different locations.

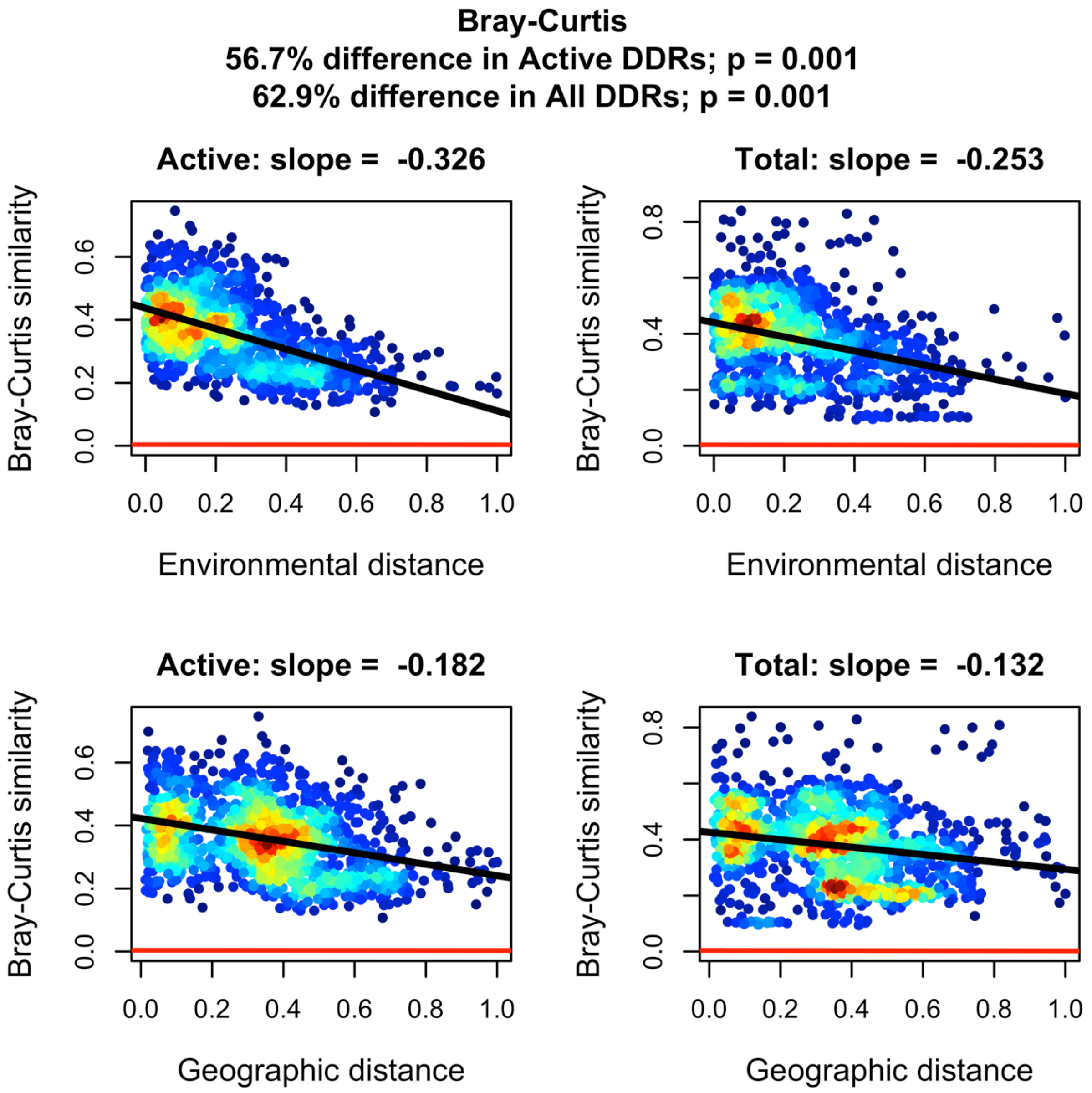

**Figure S10**. Results for distance decay relationships (DDR) for Canberra's similarity. For the active community (RNA), environmental DDRs and geographical DDRs differed by 61.7 % ($P$ = 0.001). For the total community (DNA), environmental DDRs and geographical DDRs differed by 57.8 % ($P$= 0.002). The red line is the DDR resulting from a null model whereby communities were randomly assigned to different locations.

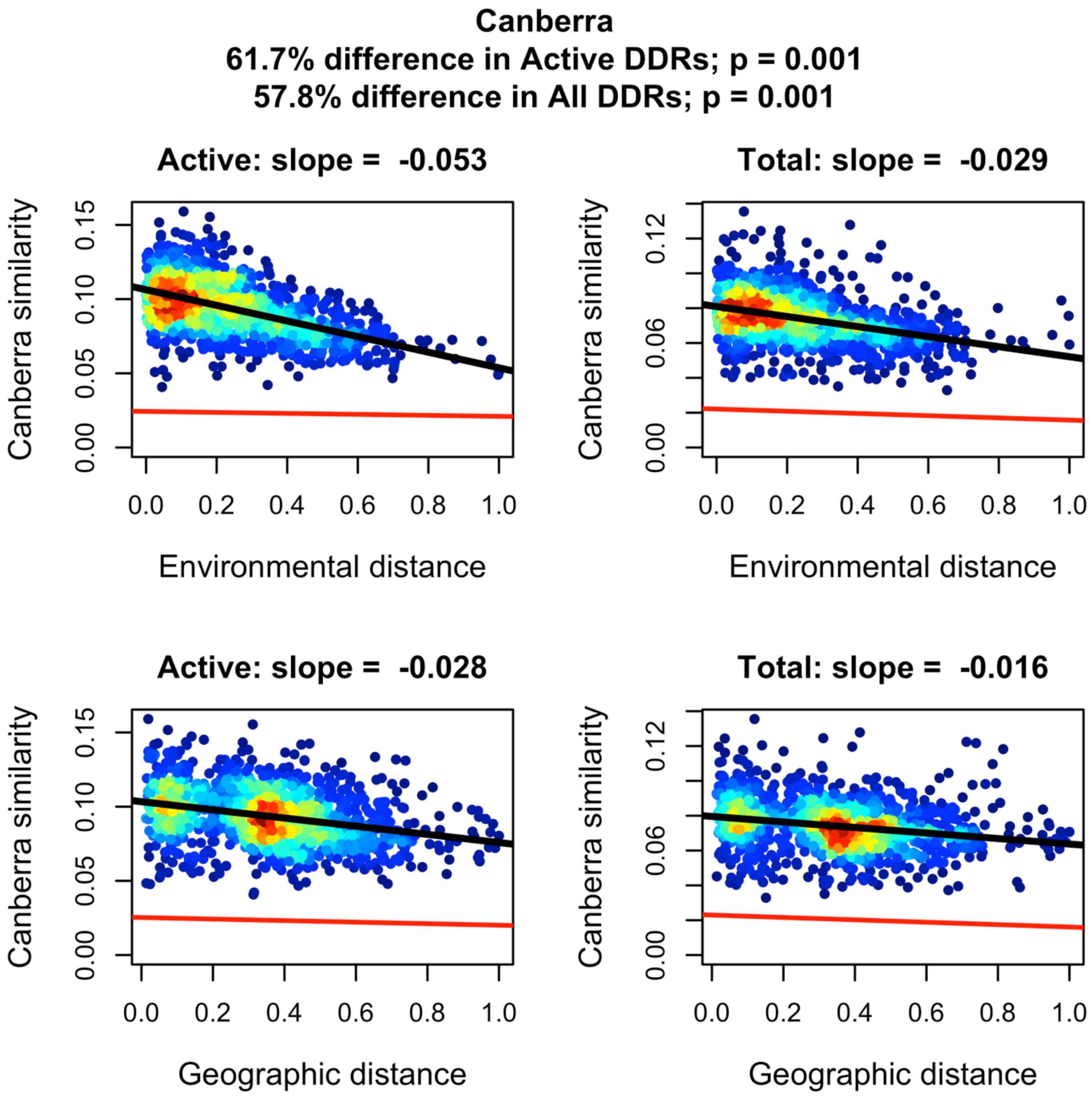

**Figure S11.** Simulation results for Sørensen's similarity. For models that met all 51 requirements of qualitatively approximating our empirical findings (Table S1), decreased death in dormancy and a greater degree of environmental filtering resulted in less percent error between models and empirical results. Neither active nor dormant dispersal had an effect on the degree to which models approximated empirical results.

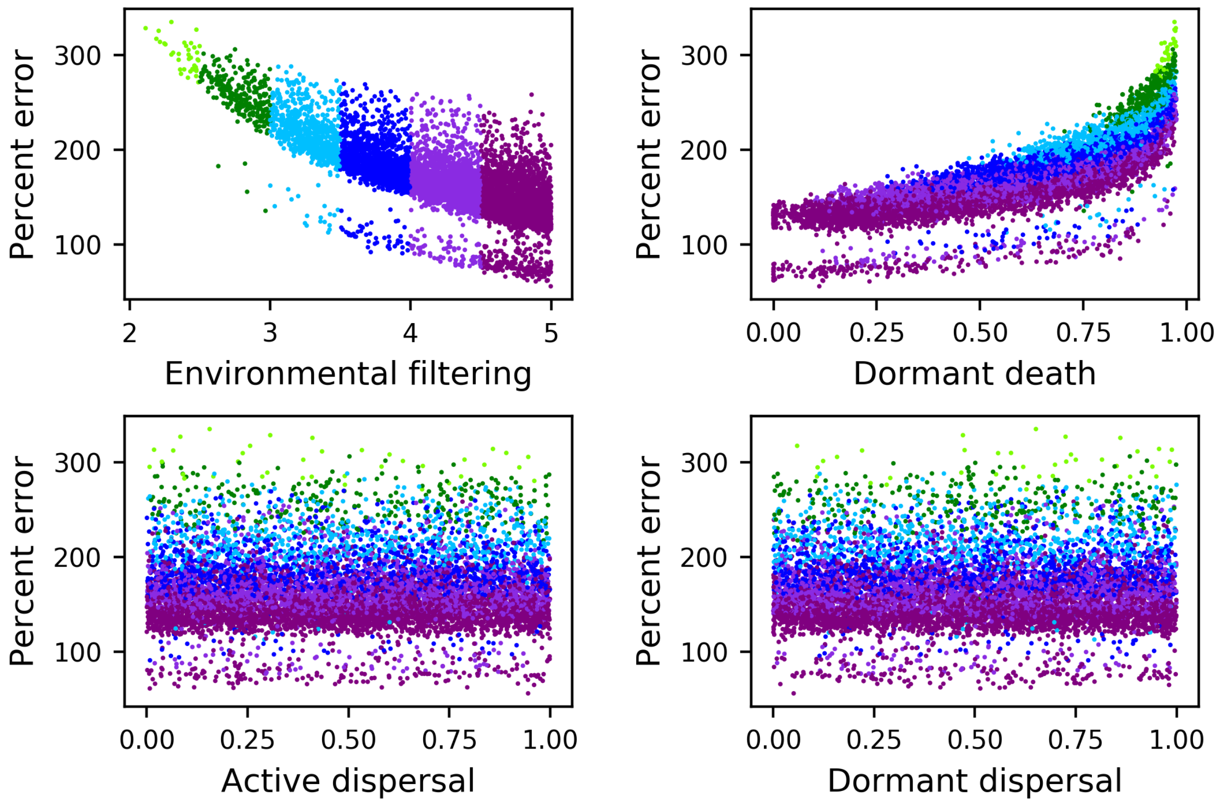

**Figure S12.** Simulation results for Bray-Curtis similarity. For models that met all 51 requirements of qualitatively approximating our empirical findings (Table S1), decreased death in dormancy and a greater degree of environmental filtering resulted in less percent error between models and empirical results. Neither active nor dormant dispersal had an effect on the degree to which models approximated empirical results.

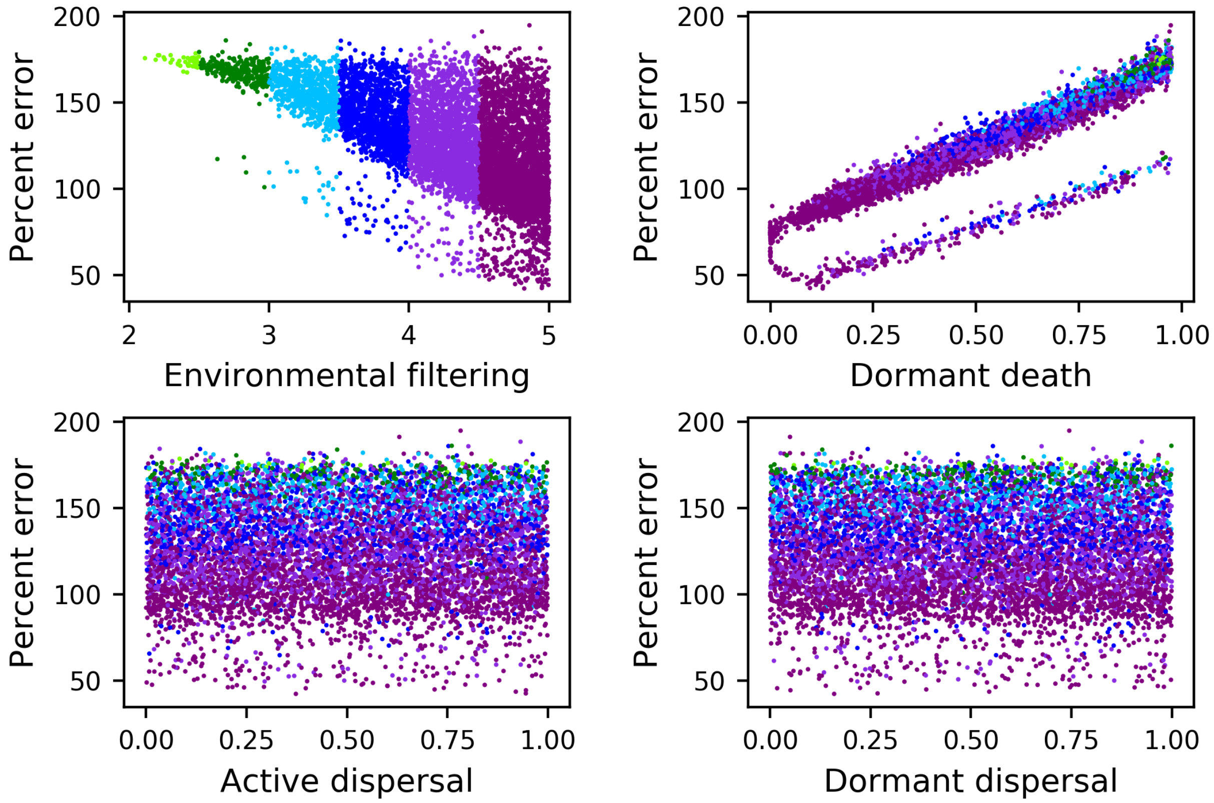

Table S1. To be counted as an appropriate simulation for the purpose of comparison to empirical results, individual simulations had to reproduce 52 aspects of our empirical DDR analyses for the active (RNA) community and the total (DNA) community. Simulations that met 50 or less of these criteria were not counted as appropriate simulations against which empirical results should be compared. Additional analyses revealed that each criterion could be met or fail to be met independent of all others. Numbers in parentheses indicate the number of times a given qualifier had to be satisfied, i.e., for each of three metrics and for multiple DDR types (e.g., environmental, geographical, RNA, DNA).

| Qualifier | Outcome |
|---|---|
| Distance-decay relationships are significant (*n* = 12) | yes/no |
| Distance-decay slopes are less than 0.0 (*n* = 12) | yes/no |
| Slopes of environmental DDRs for RNA are steeper than slopes for DNA (*n* = 3) | yes/no |
| Slopes of environmental DDRs for RNA are steeper than slopes of geographical DDRs for RNA (*n* = 3) | yes/no |
| Intercepts of environmental DDRs for RNA are greater than those for DNA (*n* = 3) | yes/no |
| Intercept of environmental DDRs for RNA are greater than intercepts of geographical DDRs for RNA (*n* = 3) | yes/no |
| Slopes of geographical DDRs for RNA are steeper than those for DNA (*n* = 3) | yes/no |
| DDR intercepts are greater than 0.0 (*n* = 12) | yes/no |
| No less than 49 communities with active and dormant populations (*n* = 1) | yes/no |
